# Mechanosensory and command contributions to the *Drosophila* grooming sequence

**DOI:** 10.1101/2023.11.19.567707

**Authors:** Shingo Yoshikawa, Paul Tang, Julie H. Simpson

## Abstract

Flies groom in response to competing mechanosensory cues in an anterior to posterior order using specific legs. From behavior screens, we identified a pair of cholinergic command-like neurons, Mago-no-Te (MGT), whose optogenetic activation elicits thoracic grooming by hind legs. Thoracic grooming is typically composed of body sweeps and leg rubs in alternation, but clonal analysis coupled with amputation experiments revealed that MGT activation only commands the body sweeps: initiation of leg rubbing requires contact between leg and thorax. With new electron microscopy (EM) connectome data for the ventral nerve cord (VNC), we uncovered a circuit-based explanation for why stimulation of posterior thoracic mechanosensory bristles initiates cleaning by the hind legs. Our previous work showed that flies weigh mechanosensory inputs across the body to select which part to groom, but we did not know why the thorax was always cleaned last. Here, the connectome for the VNC enabled us to identify a pair of GABAergic inhibitory neurons, UMGT1, that receive diverse sensory inputs and synapse onto both MGT and components of its downstream pre-motor circuits. Optogenetic activation of UMGT1 suppresses thoracic cleaning, representing a mechanism by which mechanosensory stimuli on other body parts could take precedence in the grooming hierarchy. We also mapped the pre-motor circuit downstream of MGT, including inhibitory feedback connections that may enable rhythmicity and coordination of limb movement during thoracic grooming.

## INTRODUCTION

Mechanosensation is a critical way for animals to receive information about their environment and it can trigger various behaviors from escape to grooming (Chalfie 2009, Kristan 2014). Each movement is composed of combinations of muscle activations, and more complex behavior are assembled from simpler movements selected in sequence (Palmer & Kristan 2011, Tuthill & Wilson 2016, Azevedo et al. 2020, Feng et al. 2020). A number of features makes fly grooming an excellent model system to understand how neural circuits coordinate the selection and performance of different movements (Szebenyi 1969, Seeds et al. 2014).

First, flies have to choose the correct legs to perform cleaning movements. When flies groom in response to mechanosensory cues, anterior body parts such as the eyes, antenna, and proboscis are cleaned by the front legs, while posterior parts (abdomen, wings) are cleaned by the hind legs. The notum, which is the dorsal surface of the thorax, is an exception: it can be cleaned by either front legs or hind legs, depending on the position of the bristles that are stimulated (Vandervorst & Ghysen 1980).

Second, flies must coordinate rhythmic alternation between body sweeps and leg rubbing. These movements require coordination within and between legs, using some of the same muscles in different ways. Recent work on anterior grooming, composed of head sweeps and front leg rubbing, has shown that different descending command neurons can induce either the individual movements separately or the alternating set (Guo et al. 2022).

Third, flies must select which body parts to clean if there are competing choices. When flies are dusted, they groom with an anterior to posterior progression. They usually clean the head first (eye, antenna, proboscis), then the body (abdomen, wings, and lastly thorax). There are several possible explanations for this preferred order. For example, activation of large number of sensory neurons on the head may cause stronger drive for anterior grooming (Zhang et al. 2020) or inhibitory signals may suppress posterior grooming more strongly (Seeds et al. 2014).

Recently, we and others have explored the neural bases of grooming. From neuronal activation screens with temperature-sensitive or optogenetic tools, we identified some command-like neurons that trigger grooming (Seeds et al. 2014, Hampel et al. 2015, Guo et al. 2022, Zhang et al. 2022). We also have connectome data that allows us to locate neurons of potential interest based on their connectivity to map the neural circuitry (Phelps et al. 2021, Azevedo et al. 2022).

Some of the most appealing features of grooming behavior as a model system for principles of motor control are its sequential, hierarchical organization and the rhythmic alternation of movements within its subroutines, but we have little knowledge of the neurons responsible. Here we identified a pair of command neurons, Mago-no-Te (MGT), which elicit thoracic grooming upon optogenetic activation, from a behavior screen. Then from the connectome data, we found another pair of neurons, UMGT1, that synapse onto MGT and that can inhibit thoracic grooming. Our identification of the whole sensory to motor neural circuit associated with MGT and UMGT1 gave us an entry point to understand the mechanism of some key features of grooming behavior.

## RESULTS

### Identification of Mago-no-Te (MGT)

To identify neurons involved in grooming behavior, we conducted an optogenetic activation screen using *UAS-CsChrimson* (Klapoetke et al. 2014) expressed by GAL4 and split-Gal4 lines (Pfeiffer et al. 2010) that target different neuronal populations. One split combination (*R11C07-GAL4AD* and *R45G01-GAL4DBD*) induced all posterior cleaning actions including thoracic grooming (Table S1). This line drove expression in several types of neurons (Figure S1), and we reduced that number using *hs-FLP* and *UAS-FRT-myrTopHAT2-FRT-CsChrimson-TdTomato* (Watanabe et al. 2017). Without heat-shock, basal expression of the recombinase restricted expression of CsChrimson to a single pair of neurons (Figure 1A), whose activation caused thoracic cleaning alternating with hind leg rubbing (Figure 2A-C, Video S1-3). Because of their unusual behavioral phenotype, we named these neurons “Mago-no-Te” (MGT) from the Japanese phrase “hand of the grandchild”, or less poetically “back scratcher”.

**Figure 1.**
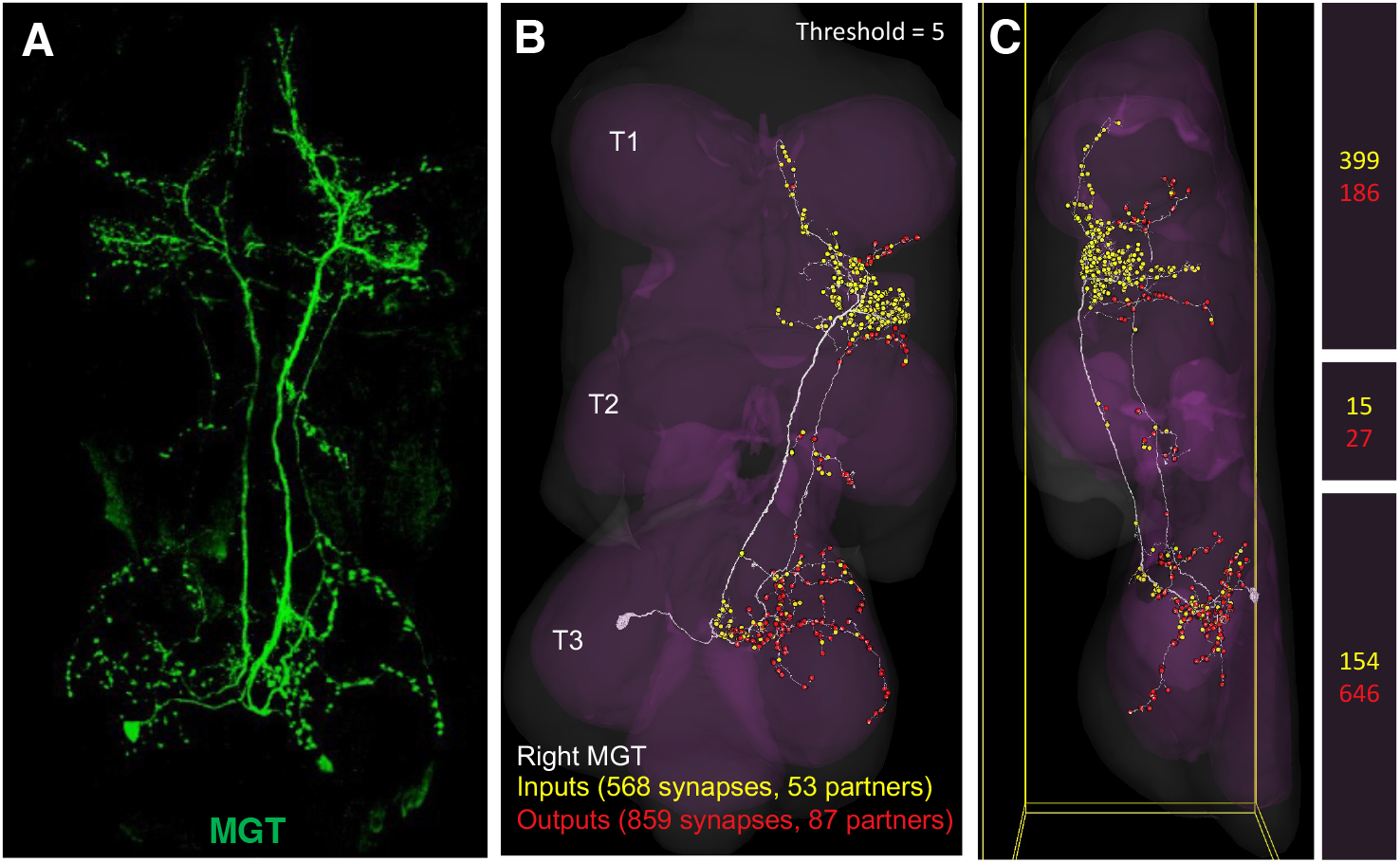
Neuroanatomy of Mago-no-Te (MGT) (A) Male VNC with MGT pair in green. Genotype: *hs-FLP; R11C07-Gal4AD, UAS-FRT-myrTopHAT2-FRT-CsChrimson-tdTomato; R45G01-Gal4DBD*. (B and C) Right MGT reconstructed from FANC EM connectome data, viewed from ventral (B) and from lateral (C) sides. More pre-synaptic inputs (yellow dots) are found between T1 and T2 thoracic segments, whereas the more post-synaptic outputs (red dots) are located in the T3. The total number of pre-synaptic inputs (yellow) and post-synaptic outputs (red) with more than five synapses per neuron in each area, based on the right side FANC dataset, is shown in the right column.

**Figure 2.**
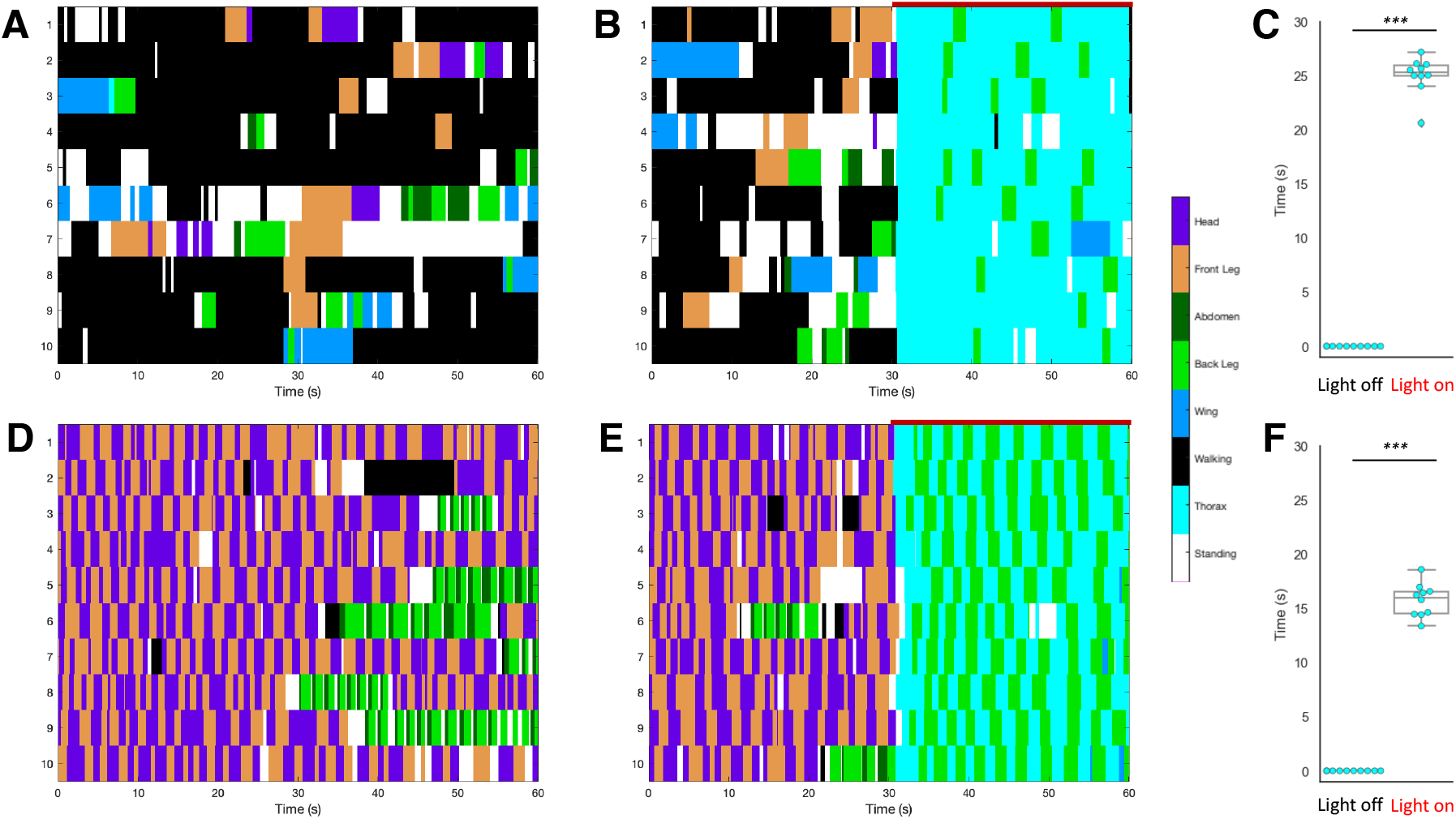
Optogenetic activation of MGT elicits thoracic grooming in both undusted and dusted flies. (A, B, D, E) Ethograms of the behavior of ten undusted (A, B) and dusted male flies (D, E) that express Chrimson in MGT. Each row shows the grooming movements of an individual fly over time. Genotype: *R11C07-Gal4AD; R45G01-Gal4DBD* split combination with *hs-FLP; UAS-FRT-myrTopHAT2-FRT-CsChrimson-tdTomato*. Color key indicates different cleaning behaviors with thoracic grooming shown in teal, scored using manual behavior annotation as described in Methods. (Activation of *MGT > Chrimson* is shown by the red line above the ethograms using 626nm at 1.12 mW/cm^2^). (C, F) The amount of total time (in seconds) that each fly (dot) spends performing thoracic grooming before (left) and during (right) light activation of neural activity in the undusted (C) and dusted (F) condition. The difference in the average time spent grooming is significant. Mann-Whitney U test was used, ***p < 0.001.

We first characterized this single pair of neurons in the VNC with light microscopic analysis (Figure 1A) and then identified them by their diagnostic morphology in electron microscopic (EM) reconstructions (Figure 1B, C). The cell body of an MGT is located in the dorsal part of the third thoracic segment (T3) and most of its pre-synaptic sites are also found in T3. Its dendrites elaborate ventrally with post-synaptic sites receiving inputs between the T1 and T2 segments (Figure 1B, C).

To determine the neurotransmitter released by MGT, we combined either *R11C07-Gal4AD* or *R45G01-GAl4AD* with Gal4DBD that label neurons based on transmitter identity. Only combinations with the cholinergic marker *ChAT[MI04508-TG4DBD]* showed thoracic grooming when optogenetically activated (Table S1). We then co-expressed MGT (*hs-FLP; R11C07-GAL4AD; R45G01-GAL4DBD > UAS-FRT-myrTopHAT2-FRT-CsChrimson-TdTomato*) with the cholinergic or glutamatergic markers, *vAchT-LexA > LexAop-StingerGFP* or *vGluT-LexA > LexAop-StingerGFP*. We found that MGT showed co-labeling only with the cholinergic marker (Figure S1C, D). Both results indicate that MGT neurons are cholinergic.

### Activation of MGT overrides dust-induced anterior grooming

Undusted flies rarely perform thoracic grooming (Figure 2A). Optogenetic activation of MGT strongly, rapidly, and repeatably induces thoracic grooming, including both the thoracic body sweeps and hind leg rubbing (Figure 2B, C). This suggests that MGT are command neurons for thoracic grooming.

Flies that are covered in dust display a hierarchical grooming sequence where head cleaning and forelimb rubbing occur first, followed by body cleaning that includes abdomen and wing sweeps alternating with back leg rubbing (Control: Figure 2D). Thoracic grooming is the last choice, or the bottom of the hierarchy, and does not happen in the first ten minutes of our assay (Seeds et al. 2014). However, dusted flies that express CsChrimson in MGT switch immediately to thoracic grooming and when optogenetically activated (Figure 2E, F). This indicates that activation of MGT can override the usual hierarchy.

### MGT are required for thoracic grooming

Since thoracic grooming is relatively rare in our standard dusting assay, silencing MGT had minimal effect. To determine whether MGT are necessary for thoracic grooming, we used a bristle deflection assay to induce thoracic grooming at a higher frequency. Stimulation of macrochaete bristles on the notum of decapitated flies elicits thoracic grooming behavior (Vandervorst and Ghysen, 1980). In our hands, 77.8% of control flies (N=18: *empty split-Gal4 > UAS-GFP-Kir2.1*) perform thoracic grooming in response to bristle deflection, while only 28.6% respond when MGT are silenced (N=14: *R11C07-GAL4AD; R45G01-GAL4DBD > UAS-GFP-Kir2.1*) (Figure S1B). This reduced response indicates that MGT are normally required for thoracic grooming.

### MGT directly command thoracic grooming but not hind leg rubbing

Optogenetic activation of MGT elicits alternation of thoracic grooming and hind leg rubbing. We hypothesized that MGT could either directly command both leg movements or that MGT could command thoracic grooming, which could in turn elicit hind leg rubbing in response to leg-thorax contact cues. To differentiate between these possibilities, we stochastically expressed CsChrimson in a single MGT, determined which side by activation of grooming, and then partially amputated either the grooming or non-grooming leg. We can still track intent to groom or rub by the movement of the remaining leg stump. We found that amputating the grooming leg (on the side where *MGT > CsChrimson* is expressed) strongly reduced the amount of time either leg spent in hind leg rubbing attempts (Figure 3A, C, Video S4, 5). In contrast, when the leg on the other side, which does not express MGT, is amputated, the stump frequently attempts hind leg rubbing following thoracic grooming sweeps by the induced, intact leg (Figure 3B, C, Video S6). These results indicate that hind leg rubbing requires successful thoracic grooming sweeps where contact occurs between leg and thorax, supporting the proposal that MGT commands thoracic grooming, which in turn induces hind leg rubbing.

**Figure 3.**
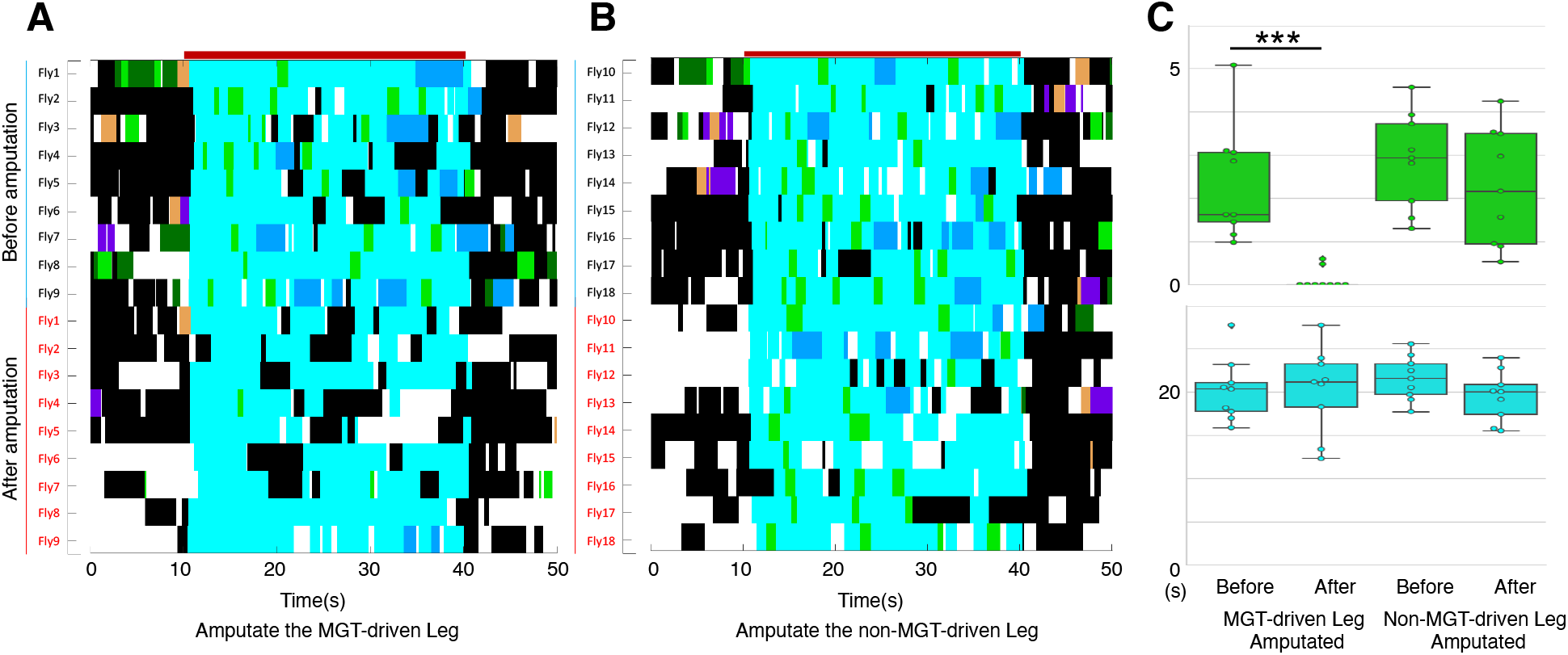
MGT activation directly commands thoracic grooming but not back leg rubbing. Ethograms of the behavior of nine flies before (top) and after (bottom) amputation of the grooming (A) and non-grooming leg (B). As in previous figures, teal color indicates thoracic grooming and lime green is hind leg rubbing; these movements can be scored in both intact legs and in the stumps of partially-amputated legs; see associated videos. Red lines indicate optogenetic activation of the MGT. Stochastic, single-sided expression in only one of the two MGT neurons is achieved by heat-shock of the following genotype (*hs-FLP; R11C07-Gal4AD, UAS-FRT-myrTopHAT2-FRT-CsChrimson-tdTomato; R45G01-Gal4DBD*). The expressing side is determined by behavioral response to light and confirmed with immunohistochemistry. (C) Box-plots showing seconds of back leg rubbing (top) and thoracic grooming (bottom) during optogenetic activation of MGT. Note that back leg rubbing is strongly reduced after the MGT-expressing leg that attempts but cannot complete thoracic grooming is amputated. Paired t-test was used. *p < 0.05, **p < 0.01, ***p < 0.001.

### Posterior macrochaete mechanosensory neurons synapse onto MGT

Previous reports indicate that forelegs will sweep the notum when anterior bristles are stimulated, while the hind legs respond to posterior bristle deflections (Vandervorst and Ghysen, 1980). Activation of MGT induces thoracic grooming exclusively with the hind legs. To determine which mechanosensory bristle neurons might connect to MGT, we expressed a green fluorescence reporter in all mechanosensory neurons (*R38B08-LexA > LexAop-mCD8-GFP*) and labeled MGT in red by expressing tdTomato. Axons of thoracic mechanosensory neurons which enter the neuropil between the first two thoracic segments T1 and T2 project in close proximity to the dendrites of MGT (Figure S2A, B). Synaptically-targeted reconstitution of fluorescence (t-GRASP) (Shearin et al. 2018) supports direct contact between mechanosensory neurons and MGT: expression of pre-t-GRASP in all mechanosensory neurons (including the thoracic bristle mechanosensory neurons) plus the expression of post-t-GRASP in MGT produced fluorescence that overlapped with the location of MGT’s dendrites (Figure S2C).

Since these experiments do not indicate which mechanosensory neurons synapse onto MGT, we turned to the VNC EM connectome data (Phelps et al. 2021, Azevedo et al. 2022). We identified thirteen pair of macrochaete mechanosensory bristle neurons, then examined the synapse number between MGT and each of the macrochaete neurons. Two of the most anterior lateral macrochaetes, uh and lh, showed few synapses. In contrast, the posterior macrochaetes such as pdc, asc or psc showed more synapses (Figure 4A-C). These results suggest that MGT elicit thoracic grooming with the hind legs because they receive more mechanosensory synapses from posterior macrochaetes.

**Figure 4.**
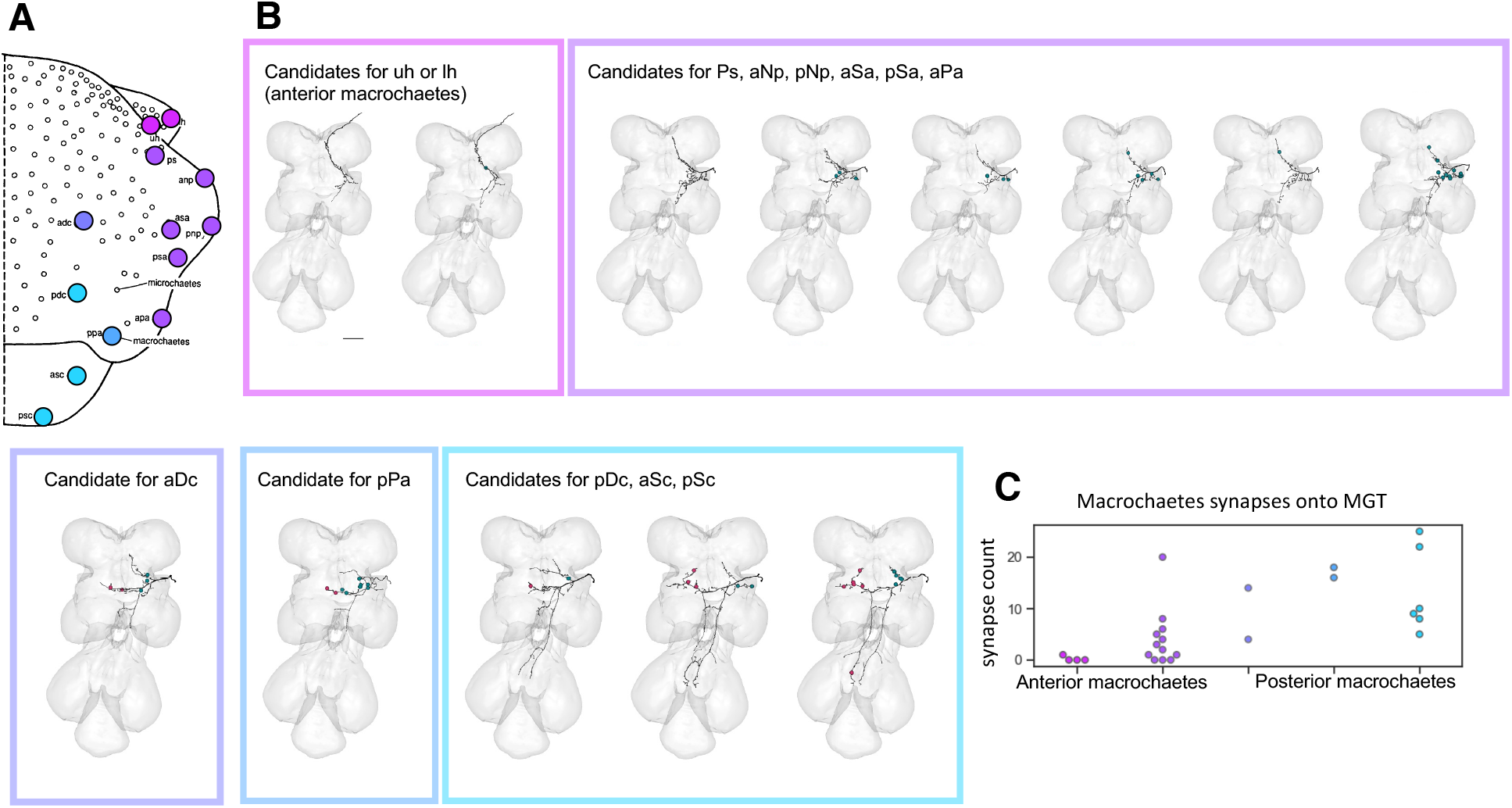
Mechanosensory neurons from posterior macrochaetes make more synaptic connection onto MGT than anterior ones. (A) Position of macrochaetes mechanosensory bristles on the notum, a schematic modified from Hartenstein and Posakony (1989). The color indicates the subgroup of macrochaetes shown in (B) and (C). Identification and proofreading of automatically-segmented neurons in the FANC EM data show axons of individual mechanosensory neurons innervating each of the 13 large thoracic bristles (macrochaetes) as they project into the VNC (B). While the identity of individual neurons cannot always be unambiguously resolved, the general class/location can be and is indicated by the colored boxes: Top left (purple) represents either uh or lh, top middle (lavender) represents ps, anp, asa, pnp, psa, apa; top right is adc, bottom left is ppa, bottom right represent either pdc, asc or psc. The dots on the EM reconstructions represent places where the mechanosensory neurons synapse onto the right and left MGT neurons (not shown). adc, ppa, pdc, asc, psc connect to both the ipsilateral and contralateral MGT, whereas the rest connects only ipsilateral; synapses with the ipsilateral partner are shown in blue-green and contra-lateral side in magenta. (C) Synapse counts of anterior and posterior macrochaetes onto both left and right MGT neurons. Since individual mechanosensory bristle neuron assignment is uncertain, the synapses are grouped by category, matching the images in the colored boxes shown in (A). Synapses were automatically identified and assigned to neurons in FANC but confirmed by manual inspection and corroborated in MANC.

### Inhibitory neurons suppress MGT

When flies groom, there is a hierarchical order of actions and thoracic grooming is always at the bottom (Seeds et al. 2014). To explain why thoracic grooming happens last, we looked for inhibitory neurons that could potentially suppress activation of MGT. From the EM connectome data, we found that the pair of neurons that are the most highly connected to MGT, pre-synaptic with the largest synapse count, are predicted to be inhibitory GABAergic from the automatic prediction of neurotransmitter identity (Eckstein et al. 2020). We named these inhibitory neurons UMGT1 (Figure 5A). UMGT1 not only synapse onto MGT but also connect to premotor and motor neurons downstream of MGT (Figure S4A). We used Maximum Intensity Projection (MIP) search (Otsuna et al. 2018) to identify split-Gal4 lines *VT031392-Gal4AD; Gad1[MI09277-TG4DBD]* that label them (Figure 5B). We then optogenetically activated UMGT1 using *UAS-CsChrimson* in a decapitated fly and stimulated bristles on the notum. Activation of UMGT1 strongly suppressed mechanically stimulated thoracic grooming behavior. While 91.3% of the control flies (N=23: *empty split-Gal4* > *UAS-CsChrimson*) demonstrate thoracic grooming in response to bristle deflection, 0% of flies where UMGT1 is activated do (N=21: *VT031392-Gal4AD; Gad1[MI09277-TG4DBD] > UAS-CsChrimson*). To verify that this is not caused by motor impairment or sluggishness, we showed that the *UMGT1 > CsChrimson* flies still respond to front leg stimulation at near normal rates (95.2% control vs. 88.9 % UMGT1 activated: Figure 5C). We also optogenetically activated intact dusted flies expressing CsChrimson in UMGT1 and found that all posterior grooming actions were reduced while anterior ones were unaffected (Figure S3A-C).

**Figure 5.**
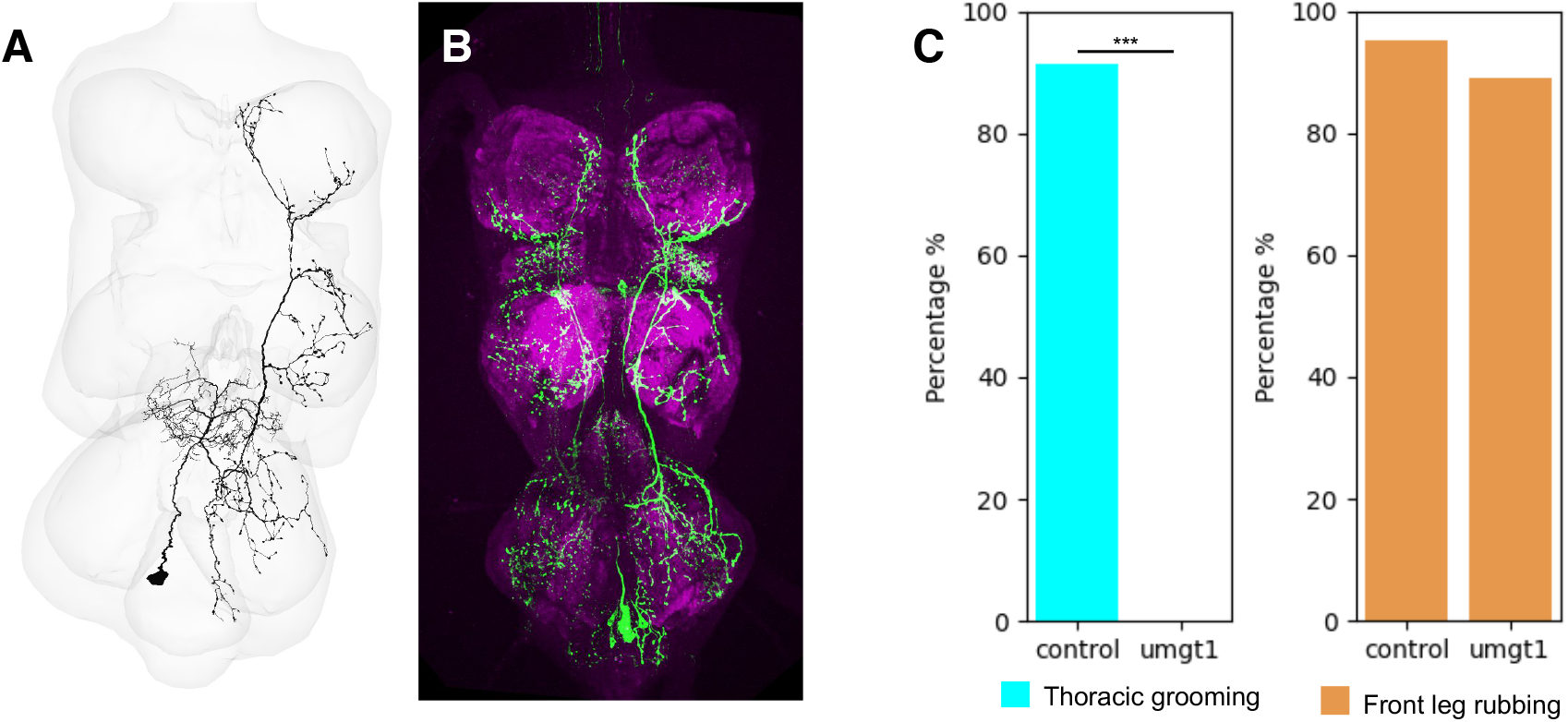
Identification of UMGT, a pair of inhibitory interneurons upstream of MGT whose activation suppresses thoracic grooming. (A) EM reconstruction of UMGT1 upstream of right MGT, ventral view of VNC. (The contralateral homolog of UMGT1 shows similar connectivity onto left MGT; data not shown.) (B) Confocal image of male VNC with genetic targeting of UMGT1 using *VT031392-Gal4AD, Gad1[MI09277-TG4DBD]* split combination driving the expression of *UAS-CsChrimson-mVenus*, stained with anti-GFP (green) and nc82 counterstain (magenta). (C) The percentage of flies that respond to bristle stimulation with thoracic grooming under control (left, N=23 flies) and constant light conditions (right, N=21 flies). Flies of this genotype do not show motor impairment and continue to respond to leg bristle stimulation with front leg rubbing under the same activation conditions, shown with orange bars in the box plot (N = 21 flies for control and N = 18 for UMGT1 > Chrimson). Genotypes are *Empty-Gal4AD, Empty-Gal4DBD > UAS-CsChrimson-mVenus* for the control and *VT031392-Gal4AD; Gad1[MI09277-TG4DBD] > UAS-CsChrimson-mVenus* for UMGT1. Chi square test of independence was used. ***p < 0.001.

We then looked for the upstream neurons of UMGT1 in the EM connectome and found haltere and wing sensory neurons directly presynaptic to UMGT1 (Figure S4A-E, Table S2). We also found contacts from other inhibitory neurons (hemi-lineage 6B and 3B), but the function of these potential dis-inhibitory motifs is still unknown. These behavioral and connectivity results indicate that UMGT1 may be key component controlling the order of grooming.

### Premotor Neuron Circuits Downstream of MGT

Since MGT activation initiates thoracic grooming by the hind legs, and most of MGT’s synapses are located in the motor region of the T3 neuropil, we investigated whether MGT targets hind leg motor neurons directly or indirectly. Using the FANC and MANC connectomes (Phelps et al. 2021, Azevedo et al. 2022, Takemura et al. 2023, Marin et al. 2023, Cheong et al. 2023), we identified the most highly connected post-synaptic partners of MGT and the shortest paths to known leg motor neurons (Figure 6A). No significant direct connections were found between MGT and T3 motor neurons. We identified three layers of highly-connected pre-motor neurons downstream of MGT. The pre-motor neuron circuits are composed of 12 single neurons or groups of neurons with similar morphology and connectivity. The first layer, direct downstream partners of MGT, is made up exclusively of excitatory cholinergic pre-motor neurons DMGT1 and DMGT Pre-motor neuron family (DMGT PMF) (Figure S5). DMGT1, the most-connected downstream partner of MGT, is a pair of T3 4B hemi-lineage local interneurons. DMGT PMF is a group of T3 contralateral homologs from the 3A hemi-lineage and receives fewer synapse connections compare to DMGT1. The secondary layer of the pre-motor network is made up of 3 groups of excitatory cholinergic neurons (DDMGT2, DDMGT PMF1, DDMGT PMF2) and an inhibitory GABAergic neuron pair (DDMGT1). The third layer is made up of glutamatergic neurons (DDDMGT2_1, DDDMGT2_3, DDDMGT PMF1_1, DDDMGT PMF2_1) and inhibitory GABAergic neurons (DDDMGT2_2, DDDMGT PMF1_2). Since glutamate can act as either excitatory or inhibitory neurotransmitter in the fly VNC, we did not assign a predicted effect to the glutamatergic neurons (Liu and Wilson 2013, Bykhovskaia and Vasin 2017). Interestingly, while layers 1 and 2 are exclusively excitatory and feedforward, DDDMGT2_2 GABAergic neurons in layer 3 provide inhibitory feedback to upper layers that could give rise to rhythmic activity.

**Figure 6.**
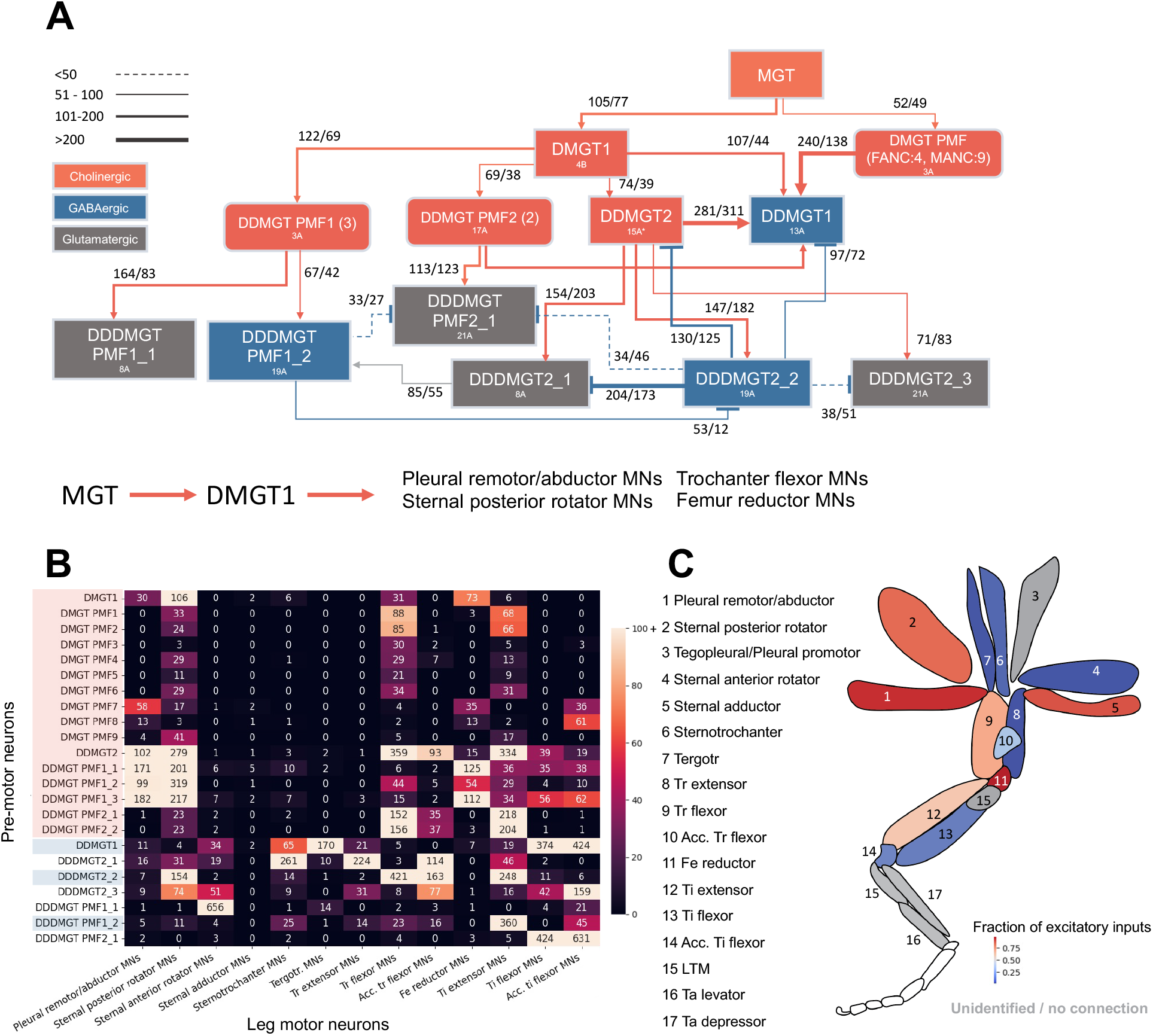
A multi-layer circuit containing both excitatory and inhibitory neurons connects MGT to leg motor neurons. (A) Schematic of pre-motor neuron circuits downstream of MGT based on FANC and MANC connectomes showing feedforward and feedback connections. Synapse counts are from FANC, while neurotransmitter and lineage information are from MANC annotation. Cholinergic (red), GABAergic (blue), and glutamatergic (grey) neurons with connection strength indicated by connection line thickness (synapse numbers of right and MGT circuits in FANC are used: right/left). “PMF” stands for pre-motor neuron family, which are a group of neurons with similar morphology and connectivity. (B) Heat map of the connectivity between the pre-motor neurons and T3 (hind leg) motor neurons. Numbers represent the synapse counts in MANC. Pre-motor neurons on the y-axis are grouped as cholinergic (red), GABAergic (blue), and glutamatergic (no color), while the motor neurons along the x-axis are in proximal to distal order as shown in (C). (C) Schematic of leg muscles targeted by MGT pre-motor neurons, modified from Cheong et. al. (2023). The color of each muscle represents the fraction of excitatory inputs from the MGT pre-motor neuron circuits.

Utilizing the MANC dataset, which presents detailed annotation for motor neurons and local interneurons in the T3 segment, we mapped which motor neurons the MGT pre-motor circuits contact (Figure 6B). To address how the pre-motor circuits are innervating leg muscles, we calculated the percentage of excitatory inputs for each identified type of leg motor neurons (Figure 6C). We found sternal posterior rotator motor neurons receive mainly excitatory inputs (83%), while sternal anterior rotator motor neurons receive significantly fewer excitatory inputs (2.4%) from the pre-motor circuits. This difference in the connectivity between body wall muscles motor neurons supports the stereotypic initiation phase of the thoracic grooming when a fly rotates their hind leg posteriorly to an extreme degree to reach the thorax. Furthermore, we found that the motor neurons innervating the tibia (Ti) receive intermediate amounts of excitatory inputs (Ti extensor: 61%, Ti flexor: 14%, Accessory (Acc) Ti flexor: 15%), which may explain the fast body sweep during thoracic grooming.

In summary, MGT indirectly connects to T3 leg motor neurons through 3 layers of pre-motor neurons, which could give rise to rhythmic activity through feedback inhibition. The connectivity between the MGT pre-motor circuits and leg motor neurons supports MGT act as command neurons for thoracic grooming.

## DISCUSSION

Here, we report the identification of the MGT command-like neurons and characterize their role in thoracic grooming behavior in *Drosophila*. Using connectome data, we describe the neural circuits connected to MGT. Posterior mechanosensory bristle neurons synapse onto MGT, instantiating a somatotopic map, consistent with the induced and natural hind-leg cleaning movements targeted at the posterior regions of the notum. Another highly connected pre-synaptic partner of MGT is a pair of inhibitory neurons, UMGT1, and activation of these neurons suppresses thoracic cleaning. This represents a potential explanation for why thoracic grooming is performed last in sequence if MGT is inhibited until other body parts are clean. Our experiments with MGT also uncover a role for sensory feedback in the assembly of the body sweeps and leg rubs that constitute the thoracic cleaning subroutine: MGT commands thoracic sweeps, but contact between leg and thorax is required to recruit the alternating hind leg rubbing. Since constant optogenetic activation of MGT initiates rhythmic leg movements, the circuits capable of generating this pattern must be located between this command-like neuron and the leg motor neurons. Our connectome analysis shows several layers of pre-motor neurons, excitatory and inhibitory, with both feed-forward and feedback connections. The sets of motor neurons that are co-innervated, directly and indirectly, suggest how the leg movements required for thoracic cleaning may be coordinated, providing a guide for future functional experiments.

### Neural control of motor subroutines

During grooming, dusted flies alternate between body sweeps and leg rubbing, where they first sweep the dust of the body onto their legs, then rub their legs to remove the dust. Our previous investigation of anterior grooming showed that different descending neurons could control front leg rubbing (DNg11) or antennal grooming (aDN) separately, or evoke the alternation of both head grooming movements together (DNg12; Guo et al 2022). Since MGT activation results in both thoracic sweeps and hind leg rubbing, we wondered whether it simply commands the entire subroutine. Arguing against this, single-sided MGT activation and amputation experiments demonstrate that MGT induces the thoracic sweep, while contact between the leg and thorax is required to induce leg rubbing attempts. This suggests that sensory cues can be important for coupling alternating motor programs in grooming subroutines.

### Somatotopy and appropriate leg choice

The *Drosophila* mechanosensory bristle neurons on the notum, the dorsal surface of the thorax, have been well studied for developmental phenomena such as cell fate decisions (Simpson et al. 1999) and axon guidance (Ghysen 1980, Chen et al. 2006, Kays et al. 2014). They were critical for the electrophysiological characterization of the nompC mechanosensitive ion channel (Walker and Zuker 2000). Although we do not know the full range of behaviors the fly uses these bristles for, previous experiments in decapitated flies showed that mechanical stimulation of anterior bristles results in a reflexive sweep of the fore-legs while similar deflection of posterior bristles induces a targeted sweep of the hind legs (Vandervorst & Ghysen 1980). This indicates the presence of a somatotopic map coupling mechanosensory input to motor output, located within the VNC. Previous anatomical characterization of the mechanosensory bristle neurons shows spatial tiling of their axons (Chen et al. 2006, Kays et al. 2014). Since activation of MGT specifically induced thoracic grooming using the hind legs, we hypothesized that MGT might receive sensory information from posterior macrochaetes. Indeed, connectome analysis shows that MGT does receive direct synaptic connections from macrochaetes, and more from posterior than anterior ones. Furthermore, MGT’s axons target the pre-motor regions associated with the hind legs. This provides a circuit-based explanation for Vandervorst and Ghysen’s behavioral demonstration that a somatotopic map informs the sensory-motor coordination needed to performed targeted scratch reflexes in the fly’s thorax.

### Assembling a sequence

Analysis of flies grooming in response to dust showed that they remove the dust in sequence, beginning with the head and then cleaning the abdomen, wings, and lastly, the thorax (Seeds 2014). From genetic activation screens, we found command-like neurons that elicit different grooming subroutines (Seeds et al. 2014, Hampel et al. 2015, Guo et al. 2022, Zhang et al. 2022), suggesting that each cleaning action can be separately controlled. But the neural basis for implementing the action hierarchy that results in sequential progression was unknown. Access to MGT now provides some insights. Activation of MGT in undusted flies induces thoracic grooming, arguing for command-like function, but activation in dusted flies also triggers thoracic grooming, out-competing anterior grooming and over-riding the hierarchy. Perhaps thoracic grooming occurs last because it takes a long time for MGT to integrate sufficient sensory inputs to fire, or because its activation is suppressed when other body parts are stimulated. We identified an inhibitory neuron, UMGT1, that receives sensory input from other regions of the body and makes many synapses onto MGT. When activated, UMGT1 can suppress thoracic grooming. These data support the second model, in which thoracic grooming occurs last because its command neuron MGT - and also the pre-motor neurons it innervates - are actively inhibited during other higher priority cleaning behaviors.

One intriguing observation is that UMGT1 receives a large number of synapses from a few haltere sensory neurons. While we do not yet know the role of this connection, we speculate that it could convey behavioral context information and explain why *Drosophila* grooming is normally suppressed during flight.

### Coordinating different leg movements

Thoracic grooming requires the hind legs to move in a specific way to sweep the notum, executing a motor program involving different leg muscles acting in order. The motor neurons controlling each leg muscle have been anatomically characterized (Baek & Mann 2009, Enriquez et al. 2015, Brierly et al. 2012) and described in the recent connectome (FANC: Phelps et al. 2021, Azevedo et al. 2022). We used these data to map the circuits connecting MGT to motor neurons and identified a complex, multi-layer network of feed-forward excitatory and feed-back inhibitory connections. With the arrival of a second VNC connectome (MANC: Takemura et al. 2023, Marin et al. 2023, Cheong et al. 2023), we were able to confirm many of these connections and assess variability in synaptic weights. While modeling and experimental work remains to disentangle how these circuits produce the rhythmic and flexible leg movements we observe, the connectome analysis suggests a few key control points to prioritize for functional studies. For example, DDDMGT2_2 (Figure 6A) is predicted to be GABAergic and its connectivity indicates two feedback inhibitory loops, suggesting that it could play a role in generating a rhythmic pattern.

MGT connects to pre-motor neurons that synapse onto motor neurons affecting related muscle groups. MGT also connects to interneurons that contact pre-motor neurons, providing a more indirect pathway to distinct muscle groups, and creating potential delay lines that could support the short-time-scale, sequential muscle contractions required for specific leg movements. Our observations of hind leg movements during thoracic body sweeps indicate that it involves flexion of the trochanter, posterior rotation around the most proximal joint (closest to the body), and then extension and flexion of the tibia. This generates the inversion of the leg to contact the thorax and the anterior to posterior sweeping movement that characterize thoracic grooming. Future work will include biomechanical modeling (NeuroMechFly 2.0: Wang-Chen et al. 2023) using information from these potential motor control circuits and functional tests of effects on limb kinematics using DeepLabCut (Mathis et al. 2018).

## ACKNOWLEDGEMENTS

We thank John Thomas for critical reading of the manuscript. For the FANC EM connectome, we thank Wei-Chung Allen Lee, John Tuthill, Jasper Phelps, and Brandon Mark. For automatic segmentation, we thank Zetta AI LLC (Ran Lu, Nico Kemnitz, Kisuk Lee, Akhilesh Halageri, Manuel Castro, Dodam Ih, Sven Dorkenwald, Forrest Collman, Casey Schneider-Mizell, Derrick Brittan, Chris S. Jordan, Thomas Macrina). For proofreader training and testing, we thank Jasper Phelps and Leila Elabaddy. We thank Ben Gorko for the initial identification of MGT in the EM dataset, and the UCSB undergraduate EM team, led by Natalie Hill and Graci Novack, for proof-reading: Anna Maciukiewicz, Elias Strategos, Ashley Mansour, Maika Grospe, Vikram Samra, Grace Ryu, Taylor Cooke, Serina Wu, Clare Andriola, Eugene Sakai, Yu Zhu, Rachel Ki, Jaxon Zhang, Carson Paxton, Estevan Mosqueda De Rosas, Dara Sadeghian, Faizan Miya, Jaxon Zhang, Myla Gupta, and Jackie Lichter. For fly strains, we thank David Anderson, Benjamin White, Janelia Research Campus, and the Bloomington Drosophila Stock Center, which is supported by a grant from the Office of the Director of the NIH under Award Number P40OD018537. This work was supported by National Institutes of Health grants NINDS NS110866 and Brain Initiative RFNS132900.

## AUTHOR CONTRIBUTIONS

S.Y., P.T., and J.H.S. conceptualized the study; S.Y. and P.T. performed experiments and analyzed data; S.Y. and P.T. wrote the original draft; J.H.S. edited the manuscript; and J.H.S. acquired the funding and supervised the study.

## STAR METHODS

### KEY RESOURCES TABLE

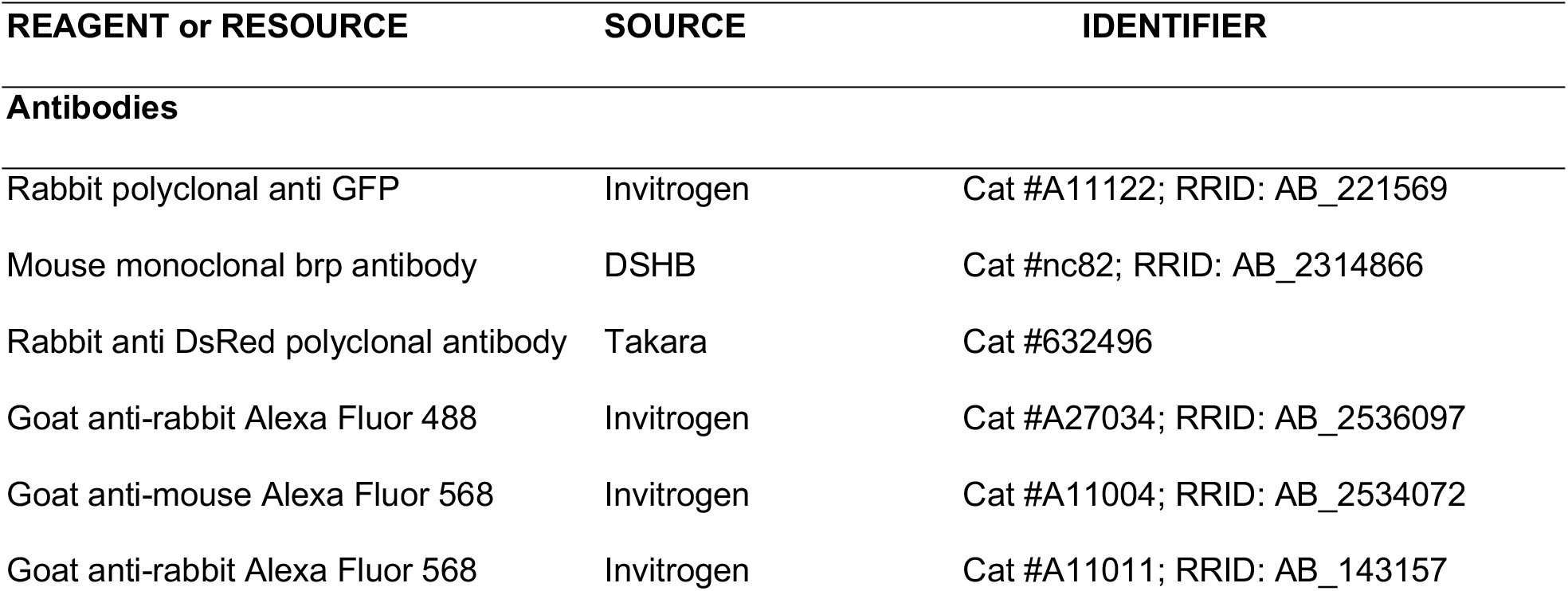
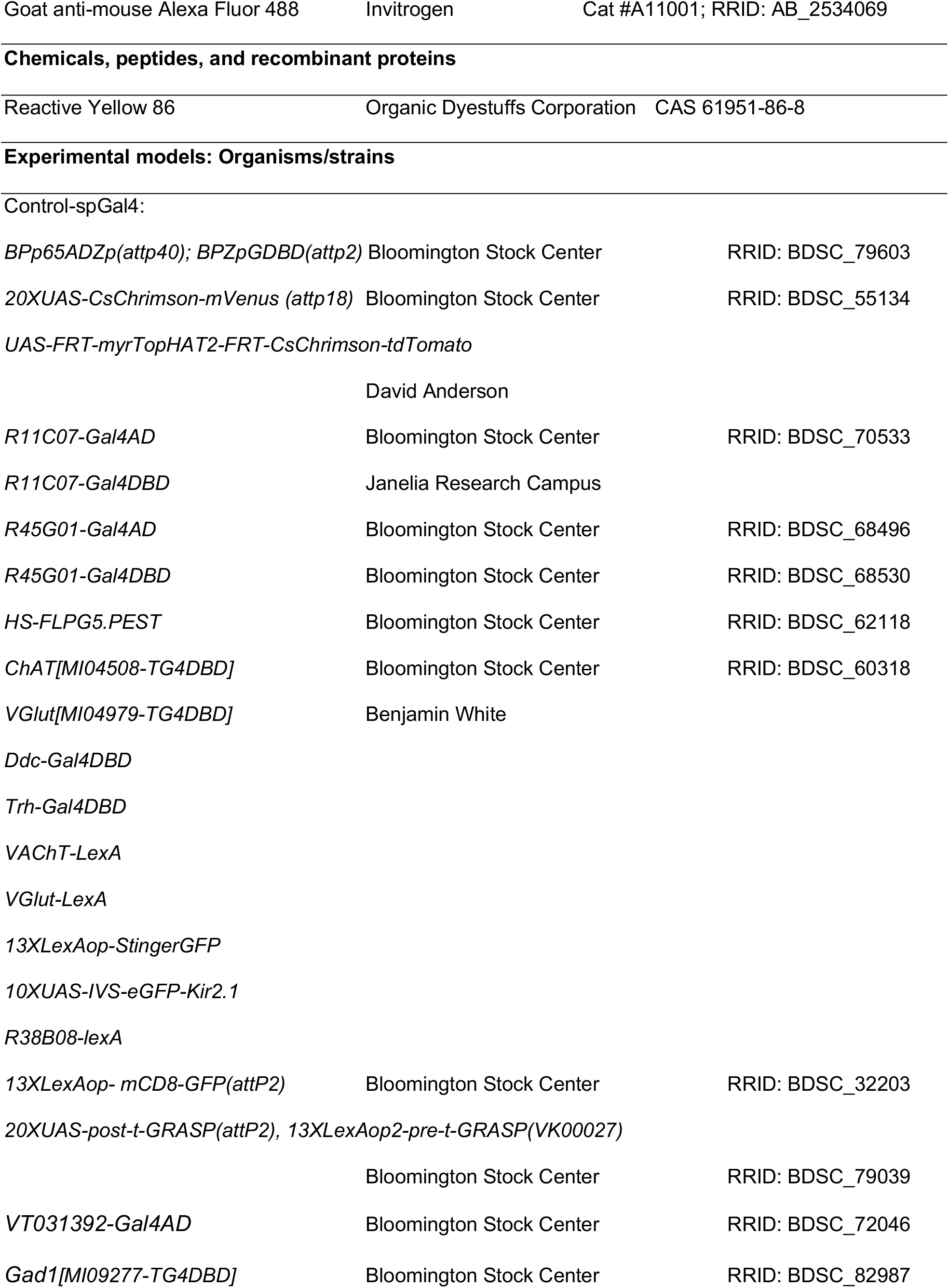
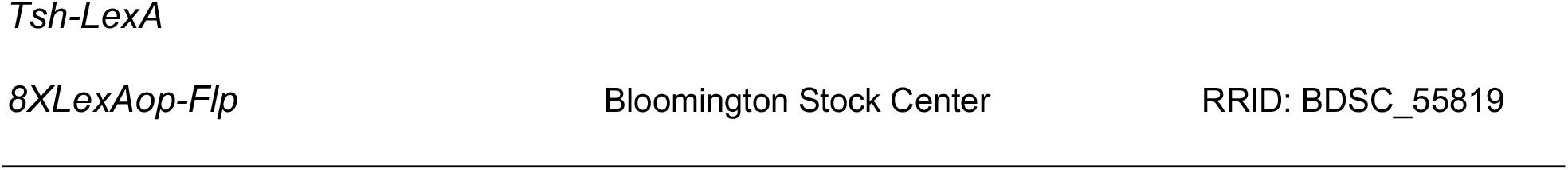

### METHOD DETAILS

#### Optogenetic experiments in undusted and dusted flies

Optogenetic activation and fly dusting were performed as described previously (Zhang et al. 2020, Guo et al. 2022) except for the amputation experiments. Custom made LED panels (LXM2-PD01-0050, 625nm) were used for light activation. Constant light with the intensity of 1.1 mW/cm^2^ was used in all experiment. 30Hz videos were recorded by IDS UI-3370CP-C-HQ camera and manually annotated in VCode.

#### Immunohistochemistry and confocal imaging

Immunohistochemistry and confocal imaging were performed as previously described (Guo et al. 2022).

#### EM reconstruction

Neuron skeletons were reconstructed in the *Drosophila* Female Adult Nerve Cord electron microscopy dataset (FANC; (Phelps et al. 2021, Azevedo et al. 2022)). MGT neuron pairs were identified based on neuroanatomical landmarks such as soma location, dendrite arborization, and backbone orientation based on the confocal images. Only 1 pair of MGT candidates was identified and extensively proofread (>4 hours) in FANC. The segment ids are 648518346503614625 for left MGT and 648518346514264286 for right MGT. A neuron is considered as “proofread” when the cell body is attached (except sensory neurons/descending neurons), the full backbone and major branches were reconstructed without attempting to add small twigs. We identified the left MGT neuron in the connectome of the *Drosophila* Male Adult Nerve Cord (MANC, Takemura et al., 2023, Cheong et al. 2023) based on the neuroanatomical features and the known connectivity pattern in FANC (segment id 21453). The MANC left MGT candidate was identified as downstream of macrochaetes (labeled as “SNta” in MANC) and UMGT1 candidates that are morphologically similar to the neurons identified in FANC, which synapse onto T3 premotor neurons that are morphologically similar to DMGT1 and DMGT PMF. We could not identify a candidate for the right MGT in MANC, and we think it likely is due to a proofreading error because we were able to identify incomplete segments resembling MGT axon projection on the other side of the VNC. For the neurons upstream and downstream of the MANC left MGT candidate, we used MANC annotations to identify the contralateral partners. We identified and proofread the 13 pairs of macrochaete thoracic mechanosensory neurons in FANC by finding the nerve where the thoracic mechanosensory neurons enter VNC. The macrochaetes were further distinguished from microchaetes and other sensory neurons in the nerve based on their axon morphology (Kays et al., 2014). EM figures are plotted from Neuroglancer interface.

#### Synaptic connectivity

The automatic synapse detection in FANC was used for studying the connectivity between neurons (Azevedo et al. 2022). We identified and proofread all the pre and postsynaptic partners of the MGT neurons with more than 5 synapses based on automatic synapse detection. UMGT1 was the major upstream of MGT and their segment ids are 648518346488618062 for the upstream of right MGT and 648518346484249895 for left MGT. We identified and proofread all the upstream partners of UMGT1 neurons with more than 5 synapses. The UMGT1 upstream of Right MGT was used for analysis. The synapse count and the FANC segment id are listed in Table S2. The hemi-lineage prediction was based on FANC and MANC. When the corresponding neuron was not annotated in both FANC and MANC, we labeled it as “unidentified”. The synapse tables were used for connectome analysis and updated on July, 2023. All the figures used the automatic synapse detection of the right MGT circuitry. The relative connection strength of right MGT and left MGT circuitry agrees.

Different criteria for proofreading were used for studying the downstream circuitry of MGT neurons because the number of predicted synapses for pre- and postsynaptic partners are high for T3 local premotor neurons. To reduce the proofreading effort, the threshold was > 10 synapses for proofreading the downstream partner of DMGT1 and the top 20 partners for all the other premotor neurons; Among the downstream partners reached the proofreading threshold, we selected the preferred partners following this criteria: 1) the neuron is a T3 local interneuron; 2) the connection agrees on both sides of the VNC; 3) the neuron can be identified in MANC based on connectivity and morphology.

We used the annotation in MANC to study the connectivity between the preferred MGT downstream partners and T3 motor neurons (Marin et al. 2023, Cheong et al. 2023). We created a connectivity matrix between preferred MGT downstream partners and T3 motor neuron types based on the annotation and automatic synapse detection in MANC.

#### Amputation experiments

A female fly with a single MGT expression was anesthetized on ice and either the grooming leg or non-grooming leg was amputated at the femur-tibia joint with a fine scissor. The fly was left for recovery in an empty vial for more than 2 hours in the dark. We used high-resolution videos (Ravbar et al. 2021) to capture the behavior from the ventral side of the fly. The side and the lid of the recording chamber were coated with insect coating and left for dry for more than 30 minutes. The amputated fly was placed into the coated recording chamber. The fly behavior was recorded with 10s light off - 30s light on - 10s light off optogenetic activation. 100Hz videos were recorded with a FLIR Blackfly S BFS-U3-13Y3M camera, and behaviors were manually annotated in VCode. For the amputated leg, we used its femur orientation and movement to characterize whether the fly is performing thoracic grooming or leg rubbing.

#### Inducing thoracic grooming with tactile stimulation

Thoracic grooming was induced via the similar method described in Kays et al. (2014). A male fly was collected and immobilized with ice. The experiment starts at 1 pm. The flies were decapitated with a fine scissor and were kept on a 1x PBS-soaked Whatman filter paper in a petri dish for 1-2 hours for recovery. The fly was used for further experiments if it was able to stand up after the recovery period. A fly was placed in a smaller petri dish under a Dual Discussion Stereo Microscope System. The video was recorded at 30 Hz with an iPhone 11 using a Microscope Smartphone Camera Adapter attached to a microscope eyepiece. Thoracic stimulation was achieved by continuous tactile stimulation of the posterior thoracic bristles with a single brush bristle for 60 seconds. The response was recorded and manually scored.

### QUANTIFICATION AND STATISTICAL ANALYSIS

Data analysis was performed in Python and MATLAB. Mann-Whitney U test was used to evaluate significance of MGT induced thoracic grooming in Figure 2. Paired t-test was used to evaluate significance in grooming behavior for optogenetic experiments in Figure 3 and Figure S3. Chi square test of independence was used to evaluate significance in the thoracic grooming and leg rubbing response in Figure S1 and Figure 5. The discrete-bout ethograms were generated from manual labels using MATLAB.

