## Supplementary data and tables for "Mechanosensory and command contributions to the *Drosophila* grooming sequence"

### Supplementary tables, figures and videos

Shingo Yoshikawa, Paul Tang, Julie H. Simpson*

Department of Molecular, Cellular, and Developmental Biology

Neuroscience Research Institute

University of California, Santa Barbara, Santa Barbara, CA 93106, USA

* Author for correspondence

### LEGENDS FOR SUPPLEMENTAL TABLES, FIGURES AND VIDEOS

**Table S1. MGT are cholinergic. Related to Figure 1 and S1.**

Different split GAL4 combinations that express *UAS-CsChrimson-mVenus* were assessed for ability to elicit thorax grooming in response to red light stimulation, indicated by +. Thoracic grooming was observed when *R11C07-GAL4AD* or *R45G01-GAL4AD*, split components that target expression to MGT, were combined with cholinergic driver *ChAT[MI04508-TG4DBD]*.

**Table S2. FANC segment ID numbers for upstream neurons of UMGT1. Related to Figure S4.**

The upstream neurons of UMGT1 with a synapse threshold of 5. The synapse table was used for connectome analysis.

**Figure S1. MGT are cholinergic and required for thoracic cleaning. Related to Figure1.**

(A) Left: the original splitGAL4 combination *R11C07-Gal4AD; R45G01-Gal4DBD* labeling MGT includes expression in additional neurons in the brain; UAS-CsChrimson-mVenus, stained with anti-GFP (green) and nc82 counterstain (magenta). Right: higher-magnification view of the VNC. (B) Silencing neural activity in MGT reduces the percentage of flies that show grooming in response to mechanical deflection of thoracic bristles. Control genotype: *Empty-Gal4AD; Empty-Gal4DBD > UAS-GFP-Kir2.1* (N=18), Experimental genotype: *R11C07-Gal4AD ; R45G01-Gal4DBD > UAS-GFP-Kir2.1* (N=14). Chi square test of independence was used. *p <0.05. (C and D) MGT expression overlaps with the cholinergic marker *vAChT-LexA* (C) but not the glutamatergic one *vGlut-LexA* (D). Male VNC with MGT (magenta) and transmitter reporter in green. Genotype: (C) *hs-FLP; R11C07-Gal4AD, UAS-FRT-myrTopHAT2-FRT-CsChrimson-tdTomato / LexAop-StingerGFP; R45G01-Gal4DBD / vAChT-LexA.* (D) *hs-FLP; R11C07-Gal4AD, UAS-FRT-myrTopHAT2-FRT-CsChrimson-tdTomato / vGlut-LexA; R45G01-Gal4DBD / LexAop-StingerGFP.*

**Figure S2. Thoracic sensory neurons synapse directly onto MGT. Related to Figure 4.**

Confocal microscopy of MGT (magenta) and mechanosensory bristle neurons (green) shows proximity in the T1/2 region of the VNC. MGT is labeled using *hs-FLP; R11C07-Gal4AD, UAS-FRT-myrTopHAT2-FRT-CsChrimson-tdTomato; R45G01-Gal4DBD*, anti-RFP (magenta). Thoracic mechanosensory neurons are targeted by *R38B08-LexA* driving *LexAop-mCD8GFP*, anti-GFP (green). The inset (B) shows the contact region at higher magnification. Note that the right front leg is amputated to eliminate those sensory projections. (C) Synaptically-targeted GFP reconstruction across synaptic partners (t-GRASP) also shows contact between mechanosensory bristle neurons and MGT. The T1/2 region of the VNC of male carrying *hs-FLP, R11C07-Gal4AD* and *R45G01-Gal4DBD* driving *UAS-FRT-myrTopHAT2-FRT-CsChrimson-tdTomato, UAS-post-t-GRASP* with *R38B08-LexA* driving *LexAop-pre-t-GRASP*, stained with anti-GFP (green) and anti-RFP (magenta). The dendrites of MGT (left) overlap with the GRASP signal (middle) between the post-synaptic sites of MGT and pre-synaptic sites of the thoracic sensory neurons, indicating that the thoracic sensory neurons directly synapse onto MGT.

**Figure S3. Activation of UMGT1 can suppress posterior grooming throughout the dust-induced grooming sequence. Related to Figure 5.**

(A) Schematic of the experimental design: flies were activated by 30-second light exposures at 2.5-minute intervals for five times (t1-t5) throughout the anterior to posterior grooming progression induced by dust. The ethograms show behaviors of eight experimental flies (B) and six control flies (C) spanning light exposure at each time point, while the histograms show total time spent performing each grooming behavior with (red) and without (blue) optogenetic activation. Genotypes: (B) *Tsh-LexA, LexAop-Flp / VT031392-Gal4AD, UAS-FRT-myrTopHAT2-FRT-CsChrimson-tdTomato; Gad1[MI09277-TG4DBD]*. (C) *Tsh-LexA, LexAop-Flp / UAS-FRT-myrTopHAT2-FRT-CsChrimson-tdTomato*. Paired t-test was used. *$p < 0.05$, **$p < 0.01$, ***$p < 0.001$.

**Figure S4. A survey of neurons pre-synaptic to UMGT1 in the FANC EM connectome reveals that haltere and wing sensory neurons are directly upstream. Related to Figure 5.**

(A) Schematic of simplified neuron circuits around UMGT1 shows the number of sensory neurons of each type and the total number of synapses they make with UMGT1. UMGT1 itself synapses onto MGT (45 synapses), as well as onto other neurons post-synaptic to MGT: the pre-motor DMGT1 neuron and three motor neurons downstream of DMGT. The numbers indicate the synapse count, automatically detected, for the right side. (B) EM reconstruction of all annotated haltere sensory neurons (SN) in the right side of VNC. Two of the four haltere sensory neurons that synapse onto UMGT1 are highlighted in cyan, with three different views, ventral, lateral and dorsal, highlighting their distinct morphology. (C, D) EM reconstruction of the ascending haltere (C) and wing (D) sensory neurons that connect onto UMGT1 with more than 5 synapses each. Different colors indicate individual neurons. (E) A survey showing all neurons pre-synaptic to the right UMGT1 connected by more than 10 synapses in rank order; number below indicates the synapses count and lineage or identity is given above where it is known. VNC is shown from the ventral view. Scale bar, 100 μm. The contralateral homolog, left UMGT1, has similar connectivity.

**Figure S5. Survey of neurons post-synaptic to MGT. Related to Figure 6.**

(A) EM reconstruction of downstream partners of right MGT with a synapse threshold of >10. Number indicates how many synapses were automatically detected between partners. Left MGT shows similar synapse number and rank orders with homologous downstream partners. View from ventral side of VNC. Scale bars, 100 μm. (B) Identification and comparison of EM reconstruction of MGT in the female (FANC: left) and male (MANC: right) datasets. Only one of the MGT neurons is identified in MANC based on morphology and up/downstream connectivity. The difference in the EM reconstruction may be due to proofreading errors.

**Table S1. MGT are cholinergic. Related to Figure 1 and S1.**

Different split GAL4 combinations that express *UAS-CsChrimson-mVenus* were assessed for ability to elicit thorax grooming in response to red light stimulation, indicated by +. Thoracic grooming was observed when *R11C07-GAL4AD* or *R45G01-GAL4AD*, split components that target expression to MGT, were combined with cholinergic driver *ChAT[MI04508-TG4DBD]*.

**Table S2. FANC segment ID numbers for upstream neurons of UMGT1. Related to Figure S4.**

The upstream neurons of UMGT1 with a synapse threshold of 5. The synapse table was used for connectome analysis.

**Video S1. Activation of MGT-induced thoracic grooming (dorsal view). Related to Figure 2.**

Genotype: *hs-FLP; R11C07-Gal4AD, UAS-FRT-myrTopHAT2-FRT-CsChrimson-tdTomato; R45G01-Gal4DBD*

**Video S2. Activation of MGT-induced thoracic grooming (ventral view). Related to Figure 2.**

Genotype: *hs-FLP; R11C07-Gal4AD, UAS-FRT-myrTopHAT2-FRT-CsChrimson-tdTomato; R45G01-Gal4DBD*

**Video S3. Activation of MGT-induced thoracic grooming (lateral view). Related to Figure 2.**

Genotype: *hs-FLP; R11C07-Gal4AD, UAS-FRT-myrTopHAT2-FRT-CsChrimson-tdTomato; R45G01-Gal4DBD*

**Video S4. Activation of one-sided MGT-induced thoracic grooming and hind leg rubbing (ventral view). Related to Figure 3.**

Genotype: *hs-FLP; R11C07-Gal4AD, UAS-FRT-myrTopHAT2-FRT-CsChrimson-tdTomato; R45G01-Gal4DBD*

**Video S5. Activation of one-sided MGT-induced thoracic grooming with partial amputation of MGT-driven leg (ventral view). Related to Figure 3.**

Genotype: *hs-FLP; R11C07-Gal4AD, UAS-FRT-myrTopHAT2-FRT-CsChrimson-tdTomato; R45G01-Gal4DBD*

**Video S6. Activation of one-sided MGT-induced thoracic grooming and leg rubbing with partial amputation of non-MGT-driven leg (ventral view). Related to Figure 3.**

Genotype: *hs-FLP; R11C07-Gal4AD, UAS-FRT-myrTopHAT2-FRT-CsChrimson-tdTomato; R45G01-Gal4DBD*

**Table S1**

| Gal4AD | Gal4DBD | Thoracic grooming |
| --- | --- | --- |
| *R11C07* | *R45G01* | + |
| *R11C07* | *ChAT[MI04508-TG4DBD]* | **+** |
| *R11C07* | *VGlut[MI04979-TG4DBD]* | **-** |
| *R11C07* | *Ddc* | - |
| *R11C07* | *Trh* | - |
| *R45G01* | *R11C07* | + |
| *R45G01* | *ChAT[MI04508-TG4DBD]* | **+** |
| *R45G01* | *VGlut[MI04979-TG4DBD]* | **-** |
| *R45G01* | *Ddc* | - |
| *R45G01* | *Trh* | - |

**Table S2**

| UMGT1 Upstream Partner | | |
|---|---|---|
| synapses count | segment id | note |
| 74 | 648518346491520424 | sensory haltere |
| 51 | 648518346506566946 | sensory haltere |
| 50 | 648518346496871578 | sensory haltere |
| 39 | 648518346524183557 | 6B |
| 37 | 648518346483818415 | 6B |
| 31 | 648518346489757711 | 6B |
| 24 | 648518346485551891 | 6B |
| 23 | 648518346476939464 | unidentified |
| 23 | 648518346467310599 | 6B |
| 21 | 648518346481767583 | 6B |
| 21 | 648518346475328312 | 3B |
| 21 | 648518346517359653 | unidentified |
| 20 | 648518346479050448 | 6B |
| 19 | 648518346518387030 | 3B |
| 19 | 648518346493957664 | sensory haltere |
| 18 | 648518346487375786 | 3B |
| 17 | 648518346480133030 | descending |
| 16 | 648518346488028190 | 6B |
| 16 | 648518346497761383 | 3B |
| 15 | 648518346483817903 | 6B |
| 15 | 648518346506609186 | 5B |
| 14 | 648518346497764455 | 1A |
| 13 | 648518346480946176 | 2A |
| 13 | 648518346485553939 | 6B |
| 13 | 648518346520339793 | 6B |
| 13 | 648518346499963135 | 6B |
| 13 | 648518346471832475 | unidentified |
| 12 | 648518346500083711 | 6B |
| 12 | 648518346470309374 | descending |

| | | |
|---|---|---|
| 11 | 648518346509919034 | 6B |
| 11 | 648518346514050841 | 5B |
| 11 | 648518346474343021 | 6B |
| 10 | 648518346502494790 | unidentified |
| 10 | 648518346483266380 | unidentified |
| 10 | 648518346495690640 | 3B |
| 10 | 648518346509023866 | descending |
| 9 | 648518346503005427 | unidentified |
| 9 | 648518346517107237 | sensory wing |
| 9 | 648518346494274951 | unidentified |
| 9 | 648518346512055999 | unidentified |
| 9 | 648518346496049164 | unidentified |
| 9 | 648518346480649985 | sensory haltere |
| 8 | 648518346524038917 | unidentified |
| 8 | 648518346487847703 | 3B |
| 8 | 648518346501911369 | sensory haltere |
| 8 | 648518346520340049 | unidentified |
| 8 | 648518346496703460 | sensory wing |
| 8 | 648518346483654575 | unidentified |
| 8 | 648518346475459681 | unidentified |
| 8 | 648518346474176109 | unidentified |
| 7 | 648518346508810559 | unidentified |
| 7 | 648518346490274314 | sensory wing |
| 7 | 648518346510843522 | sensory wing |
| 7 | 648518346479777765 | sensory wing |
| 7 | 648518346482123668 | 3B |
| 7 | 648518346494921866 | 18B |
| 7 | 648518346514303943 | unidentified |
| 7 | 648518346478033109 | sensory haltere |
| 7 | 648518346487335082 | sensory wing |
| 6 | 648518346485963154 | descending |
| 6 | 648518346507138376 | unidentified |
| 6 | 648518346486065288 | 3B |

| | | |
|---|---|---|
| 6 | 648518346481497281 | unidentified |
| 6 | 648518346476928784 | unidentified |
| 6 | 648518346472925540 | descending |
| 6 | 648518346488541880 | unidentified |
| 6 | 648518346479497408 | unidentified |
| 6 | 648518346494223495 | sensory haltere |
| 6 | 648518346476455798 | unidentified |
| 6 | 648518346496263997 | unidentified |
| 5 | 648518346513149265 | sensory wing |
| 5 | 648518346501836394 | unidentified |
| 5 | 648518346492716219 | 5B |
| 5 | 648518346509606714 | sensory wing |
| 5 | 648518346514429639 | sensory haltere |
| 5 | 648518346476900624 | unidentified |
| 5 | 648518346511769712 | unidentified |
| 5 | 648518346476467830 | 17A |
| 5 | 648518346514037017 | 7B |
| 5 | 648518346509079077 | unidentified |
| 5 | 648518346472292658 | 7B |
| 5 | 648518346496353655 | sensory haltere |
| 5 | 648518346498898753 | descending |
| 5 | 648518346467310855 | sensory wing |

**Figure S1**

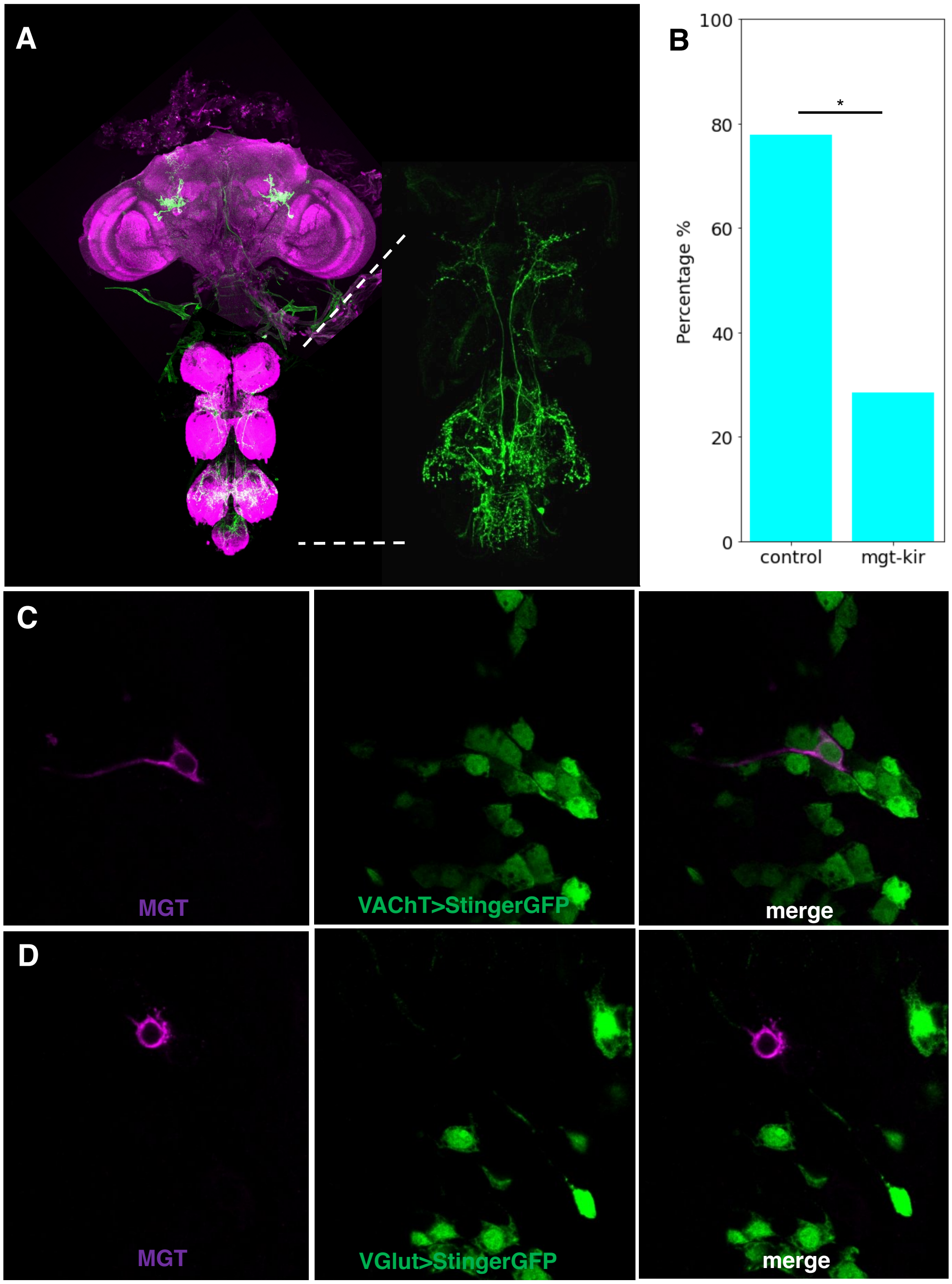
A
B
100
80
60
40
20
0
Percentage %
*
control
mgt-kir
C
MGT
VAChT>StingerGFP
merge
D
MGT
VGlut>StingerGFP
merge

**Figure S2**

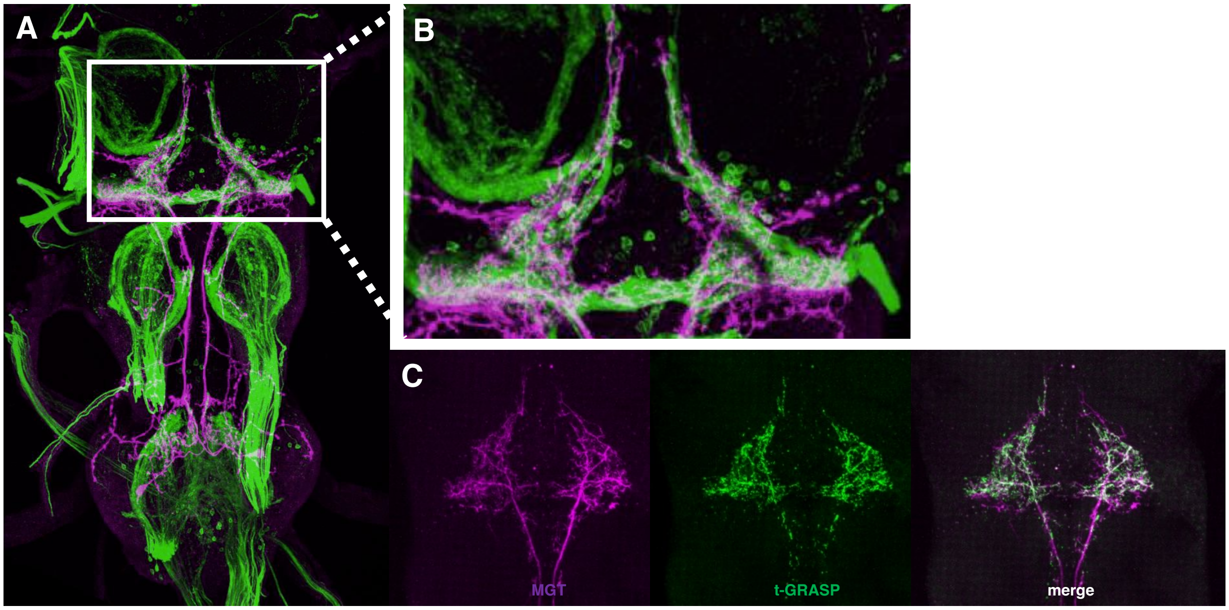

### Figure S3

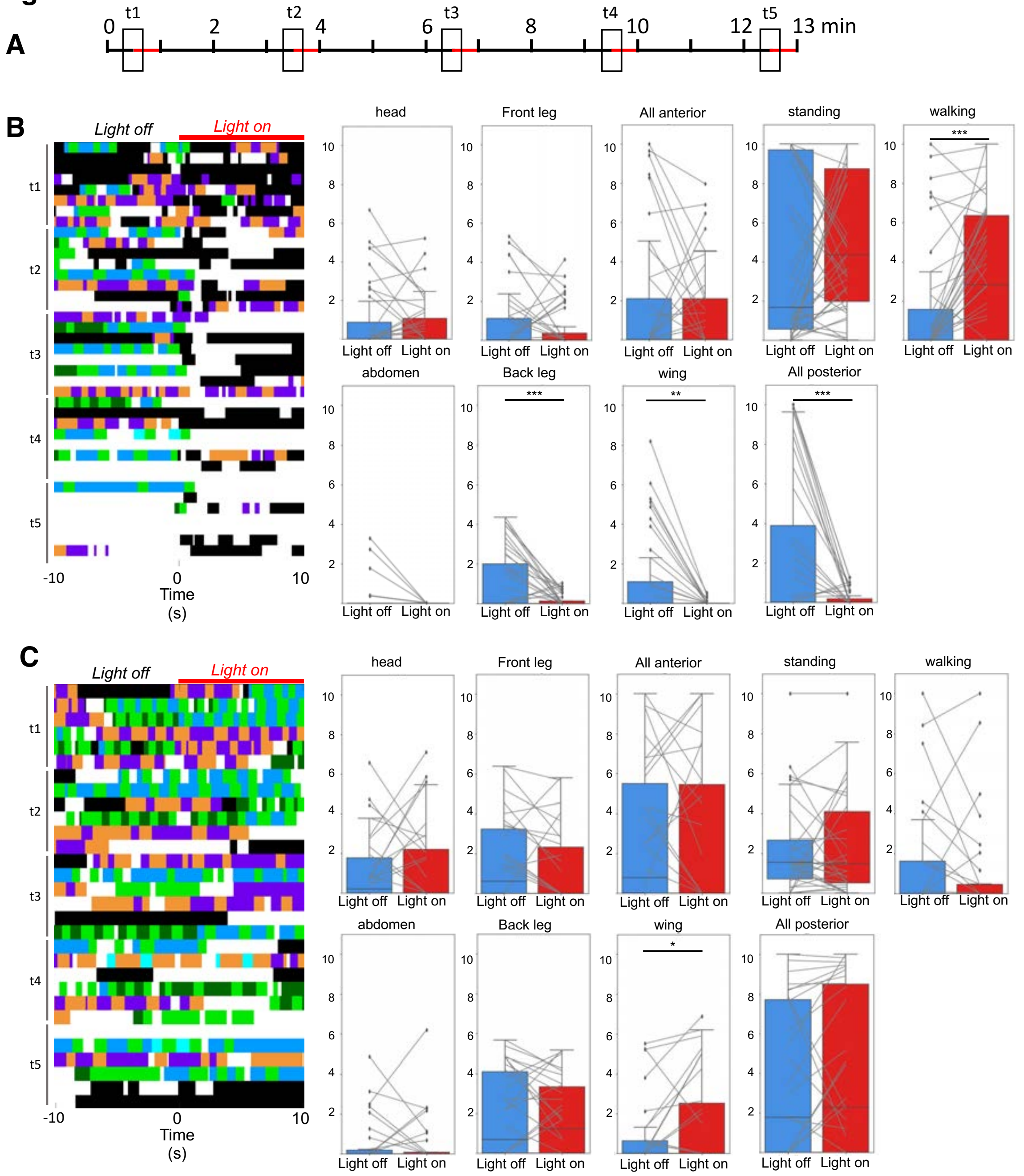
A
0
t1
2
t2
4
6
t3
8
t4
10
12
t5
13 min
B
Light off
Light on
t1
t2
t3
t4
t5
-10
0
10
Time
(s)
head
Front leg
All anterior
standing
walking
Light off
Light on
abdomen
Back leg
wing
All posterior
***
**
C
Light off
Light on
*

### Figure S4

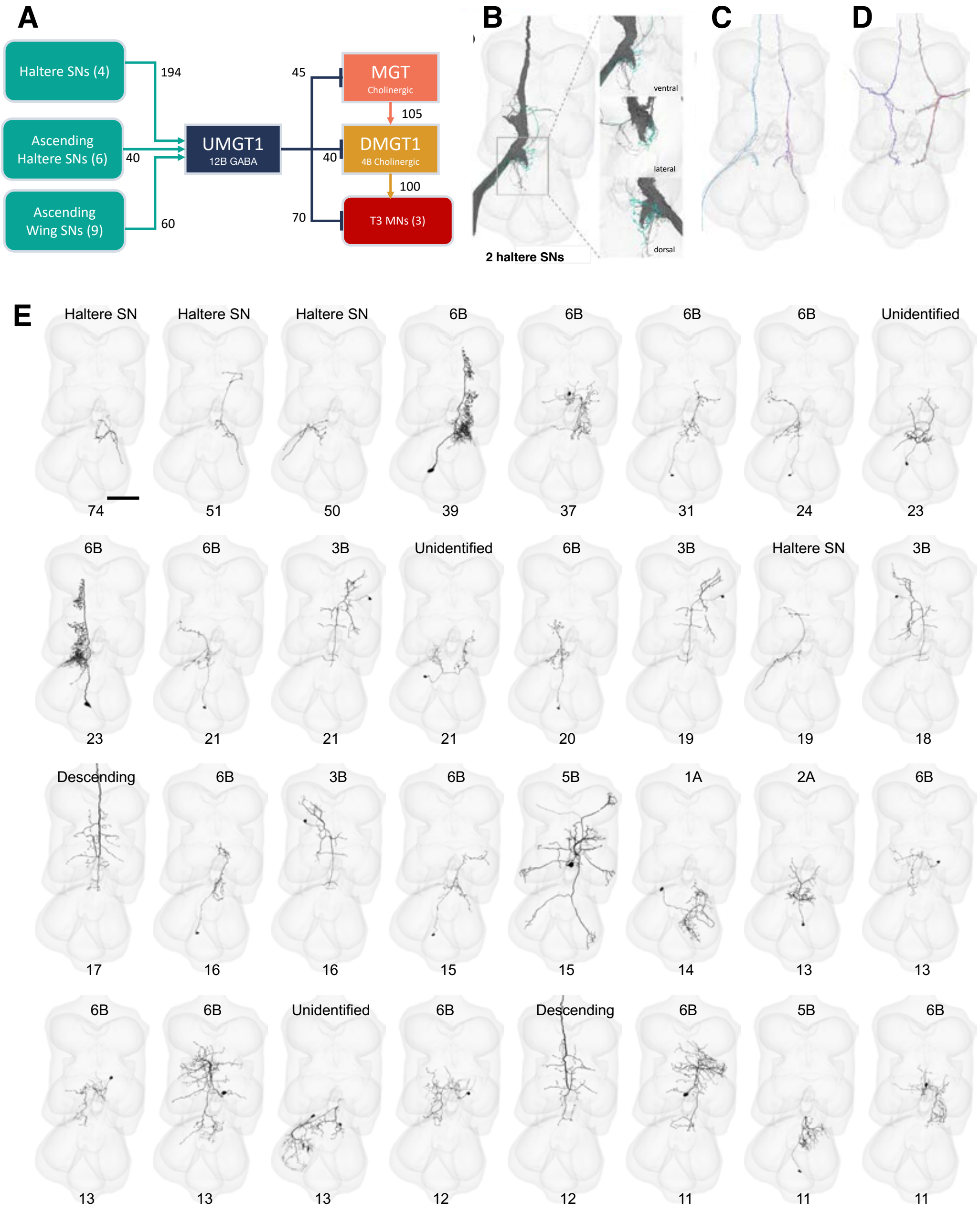
A
Haltere SNs (4)
Ascending Haltere SNs (6)
Ascending Wing SNs (9)
194
40
60
UMGT1
12B GABA
45
40
70
MGT
Cholinergic
105
DMGT1
4B Cholinergic
100
T3 MNs (3)
B
ventral
lateral
dorsal
2 haltere SNs
C
D
E
Haltere SN
Haltere SN
Haltere SN
6B
6B
6B
6B
Unidentified
74
51
50
39
37
31
24
23
6B
6B
3B
Unidentified
6B
3B
Haltere SN
3B
23
21
21
21
20
19
19
18
Descending
6B
3B
6B
5B
1A
2A
6B
17
16
16
15
15
14
13
13
6B
6B
Unidentified
6B
Descending
6B
5B
6B
13
13
13
12
12
11
11
11

### Figure S5

A

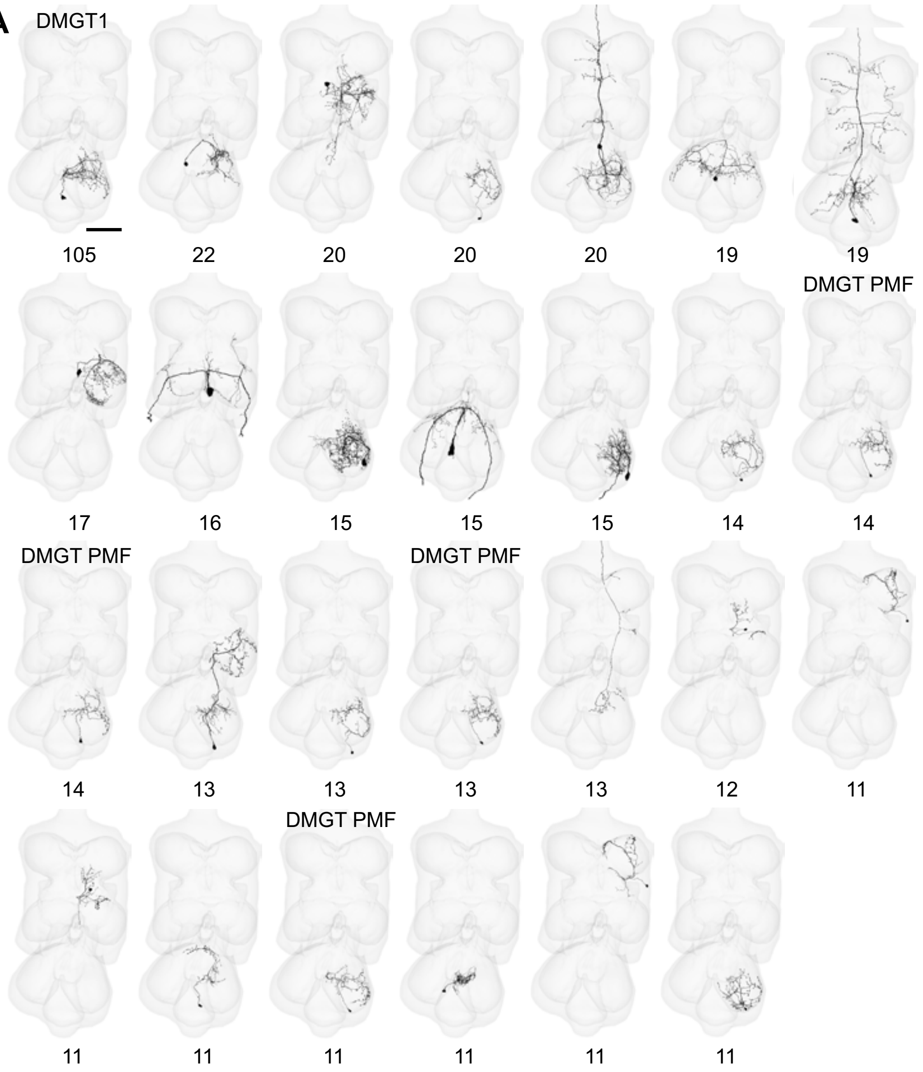
DMGT1
105
22
20
20
20
19
19
DMGT PMF
17
16
15
15
15
14
14
DMGT PMF
14
13
13
DMGT PMF
13
13
12
11
DMGT PMF
11
11
11
11
11
11

B

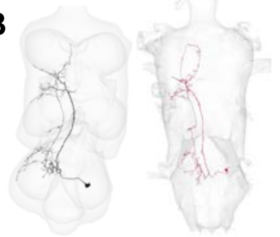
